# 1,4-dihydroxy Quininib Attenuates Growth of Colorectal Cancer Cells and Xenografts and Regulates the TIE-2 Signaling Pathway in Patient Tumours

**DOI:** 10.1101/503888

**Authors:** Clare T. Butler, Susan Kennedy, Amy Buckley, Ronan Doyle, Emer Conroy, William M. Gallagher, Jacintha O’Sullivan, Breandán N. Kennedy

**Affiliations:** UCD School of Biomolecular and Biomedical Science, UCD Conway Institute, Dublin D04 V1W8, Ireland.; Trinity Translational Medicine Institute, Department of Surgery, Trinity College Dublin, St. James’s Hospital, Dublin 8, Ireland.; Department of Histopathology, University of Dublin, Trinity College and the Central Pathology Laboratory, St James’s Hospital, Dublin 8, Ireland

**Keywords:** Colorectal Cancer, Angiogenesis, Cysteinyl Leukotriene, Drug Discovery, Xenograft, Pharmacology, metastasis, G-protein-coupled receptor (GPCR), Calpain, Vascular Endothelial Growth Factor (VEGF), Vascular Cell Adhesion Molecule 1 (VCAM-1), TIE-2

## Abstract

Colorectal cancer (CRC) is the second leading cause of cancer associated deaths in developed countries. Cancer progression and metastatic spread is reliant on new blood vasculature, or angiogenesis. Tumour-related angiogenesis is regulated by pro- and anti-angiogenic factors secreted from malignant tissue in a stepwise process. Previously we structurally modified the small anti-angiogenic molecule quininib and discovered a more potent anti-angiogenic compound 1, 4 dihydroxy quininib (Q8), an antagonist of cysteinyl leukotriene receptor-1 with VEGF-independent bioactivity. Here, Q8, quininib (Q1) and five structural analogues were assayed for anti-tumorigenic effects in pre-clinical cancer models. Q8 reduced clone formation of the human colorectal cancer cell line HT29-Luc2. Gene silencing of CysLT_1_ in HT29-Luc2 cells significantly reduced expression of calpain-2. In human *ex vivo* colorectal cancer tumour explants, Q8 significantly decreased the secretion of both TIE-2 and VCAM-1 expression. *In vivo* Q8 was well tolerated up to 50 mg/kg by Balb/C mice and significantly more effective at reducing tumour volume in colorectal tumour xenografts compared to the parent drug quininib. In tumour xenografts, Q8 significantly reduced expression of the angiogenic marker calpain-2 in. In summary, we propose Q8 may act on the TIE-2-Angiopoietin signalling pathway to significantly inhibit the process of tumour angiogenesis in colorectal cancer.

## Introduction

According to the American Cancer Society ∼40% of patients with colorectal cancer (CRC) are diagnosed with localised disease. As tumours depend on angiogenesis for growth and metastasis this biological process is focal to anti-cancer drug development (1). The anti-VEGF biological bevacizumab was FDA approved in 2004 for treatment of metastatic colorectal cancer (mCRC) in combination with Ironotecan, 5-fluorouracil and leucovorin (IFL) and in 2006 granted approval for second-line treatment of mCRC in combination with 5-fluorouracil, leucovorin and oxaliplatin (FOLFOX) (2). However, bevacizumab only extends life expectancy by approximately 4.7 months in 40% of patients and can cause gastrointestinal perforation, hypertension and bleeding (3). The development of less toxic anti-angiogenic drugs with improved efficacy is warranted.

Angiogenesis, a highly orchestrated process between endothelial cells and pericytes occurs pathologically in response to hypoxia and inflammation in the tumour microenvironment. The progression and metastatic spread of tumour cells is reliant on angiogenesis (4) and endothelial cells respond to factors secreted from the malignant cells. Through the release of growth factors and direct contact with endothelial cells, pericytes contribute to vessel maturation (6) while endothelial cells can regulate the bidirectional movement of immune cells into and out of the blood. Inducers of angiogenesis include vascular endothelial growth factors (VEGFs), angiopoietins (ANG’s), transforming growth factors (TGFs), platelet derived growth factors (PFGF), tumour necrosis factor-α (TNF-α), interleukins and fibroblast growth factor (FGF) proteins.

TIE-1 and TIE-2 (also known as TEK) are receptor tyrosine kinases for the angiopoietin growth factor ligands of which ANGPT1 and ANGPT2 are best characterized (5). TIE receptors are largely expressed in the endothelium with expression also observed in a small subset of TIE-2 expressing monocytes/macrophages (TEMs) (6). Recently, pericytes were reported to express TIE-2, albeit at a much lower levels than endothelial cells (7). In early tumour development, TIE-2 expressing TEMs are recruited to solid tumours to facilitate their vascularization and growth (8). In pericytes, TIE-2 controls the expression of Calpain1, a key regulator of cellular migration and invasion and important in tumour neovascularisation (7). The ANGPT-TIE pathway plays an important role in blood vessel development, remodelling and stability influencing angiogenesis and metastasis (9).

Chronic inflammation is a predisposing factor for the development of many cancers, particularly for colorectal cancers (10). Cysteinyl leukotrienes (CysLTs) LTC_4_, LTD_4_ and LTE_4_ are products of cell-membrane bound arachidonic acid (AA) metabolism and can induce inflammation by activating the cysteinyl leukotriene receptors CysLT_1_ and CysLT_2_ and the less functionally studied GPR99 (CyLT_3_) (14,15). Agonism of CysLT_1_ by LTD_4_ can affect cell proliferation and cell cycle regulation (10,11). The connection between angiogenesis and cysteinyl leukotriene signalling has been demonstrated using murine aortic ring models (12). Treatment of murine aortic explants with the CysLT_1_ antagonist Montelukast could block the growth of vascular sprouts when cultured in 25% serum, however, antagonism of the CysLT_2_ using BAY-cyslt2 did not prevent sprouting (12). Our parent compound Quininib, a previously developed CysLT_1_ antagonist is a known anti-angiogenic agent and inhibitor of tumour growth in xenografted mice (13-15).

In this study, we demonstrate that quininib analogues, predominantly the CysLT_1_ antagonist Q8, elicit robust anti-cancer activity in pre-clinical models and in human *ex vivo* colorectal patient tumour explants. In HT29-Luc2 CRC cells, Q8 reduces long-term proliferation, and gene silencing of CysLT_1_ is sufficient to significantly reduce calpain-2 expression. Q8 has excellent safety pharmacology when administered to mice up to 50 mg/kg. Q8 significantly reduced tumour volume in mouse colorectal tumour xenografts compared to vehicle control. Q8 significantly reduced expression of angiogenic marker calpain in tumour xenografts. In human patient CRC explants, Q8 significantly reduced the secretions of TIE-2 and VCAM-1. Overall, Q8 acts in an alternative pathway, non-redundant to the VEGF pathway, and may represent an alternative treatment strategy to counteract anti-VEGF resistance in CRC.

## Results

### Quininib analogues reduce HT29-Luc2 colony formation

To determine if structural analogues of quininib, that significantly block angiogenesis can effectively attenuate cell proliferation, colony formation assays were conducted in HT29-Luc2 colorectal cells (13). Treatment of HT29-Luc2 cells for 24, 48, 72 or 96 hours reduced average clone survival 10 days later to ∼21% (*p*<0.001) with 10 µM quininib (Q1) and ∼56% with 10 μM 5-fluorouracil (*p*<0.05) compared to ∼100% survival with 0.1% DMSO (control) (**Figure 1 A & B**). 10 µM of quininib analogues Q22 and Q18 significantly reduced average clone survival to ∼57% (*p*<0.05) and ∼27% (*p*<0.001) of control, respectively. Clone survival observed with 10 µM Q8, P4 and P18 were much greater at ∼92%, ∼106% and ∼95%, respectively. 20 μM Q1 reduced average clone survival to ∼6% compared to ∼21% with 20 μM 5-fluorouracil, both significantly reduced compared to 0.1% DMSO control (*p*<0.001). Q22 and Q18 were more cytotoxic at 20 μM, and average clone survival over 96 hours was ∼21% and ∼2%, respectively (*p*<0.001). 10 μM Q8 had no effect on clone survival but 20 μM Q8 significantly reduced average clone survival over 96 hours to ∼25% (*p*<0.001) (**Figure 1 A**). 20 μM of P18 or P4 analogues did not significantly affect clone survival. In summary, quininib (Q1), Q22 and Q18 were cytotoxic to HT29-Luc2 clones at both 10 and 20 μM. P18 and P4 were not cytotoxic to cells at 10 or 20 μM. Q8 was not cytotoxic at 10 μM but significantly reduced clone survival at 20 μM.

**Figure 1.**
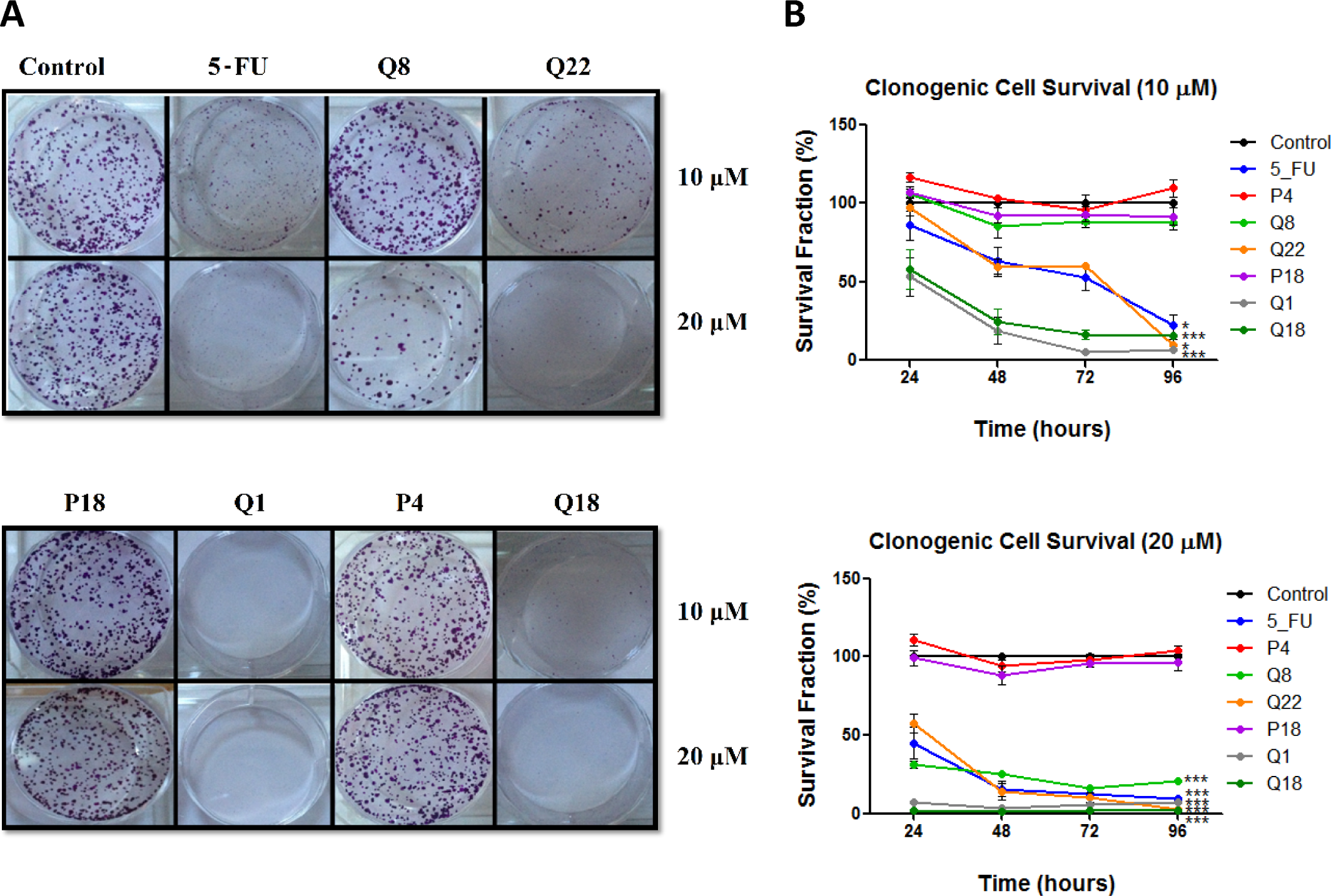
Quininib analogues reduce HT29-Luc2 colony formation. **A.** Images of clones captured by digital photography after 10 days of culture following treatment with 10 or 20µM analogues for 48 hours. Clones were stained with 0.5% crystal violet before counting. **B.** Graphs show the percentage survival fraction of clones at 24, 48, 72 and 96 hours post analogue treatment. 1,500 cells were seeded and treated in duplicate in 6-well plates for each individual experiment and individual experiments were conducted three times (n=3). Statistical analysis was performed by ANOVA with Dunnett’s post hoc multiple comparison test. Error bars are mean +S.E. *, *p*<0.05; ***, *p*<0.001.

### CysLT_1_ nuclear expression in HT29-Luc2 cells regulates downstream effectors calpain-2 and NF-ĸB

To determine if CysLT_1_, the cognate receptor for quininib and analogues, regulates calpain-2 and NF-kB in HT29-Luc2 colorectal cancer cells, immunodetection and gene silencing were applied. As in human microvascular endothelial cells (27), CysLT_1_ is abundantly expressed in the nuclear compartment of HT29-Luc2 cells but not in the cytoplasm (**Figure 2 A**). 20 nM of a 27mer siRNA significantly silenced CysLT_1_ protein expression by ∼70% compared to a scrambled siRNA control (**Figure 2 B**). 20 nM CysLT_1_ siRNA also significantly decreased calpain-2 expression compared to control (*p*=0.0268) (**Figure 2 C**). ELISA quantification of activated NF-ĸB p65 in HT29-Luc2 cells showed significant reductions (∼35%) when treated with 20 nM CysLT_1_ siRNA compared to untreated or scrambled siRNA controls (*p*<0.01) (**Figure 2 D**). In summary, CysLT_1_ silencing in HT29-Luc2 cells significantly reduced levels of downstream pro-angiogenic or pro-inflammatory proteins calpain-2 and NF-ĸB.

**Figure 2.**
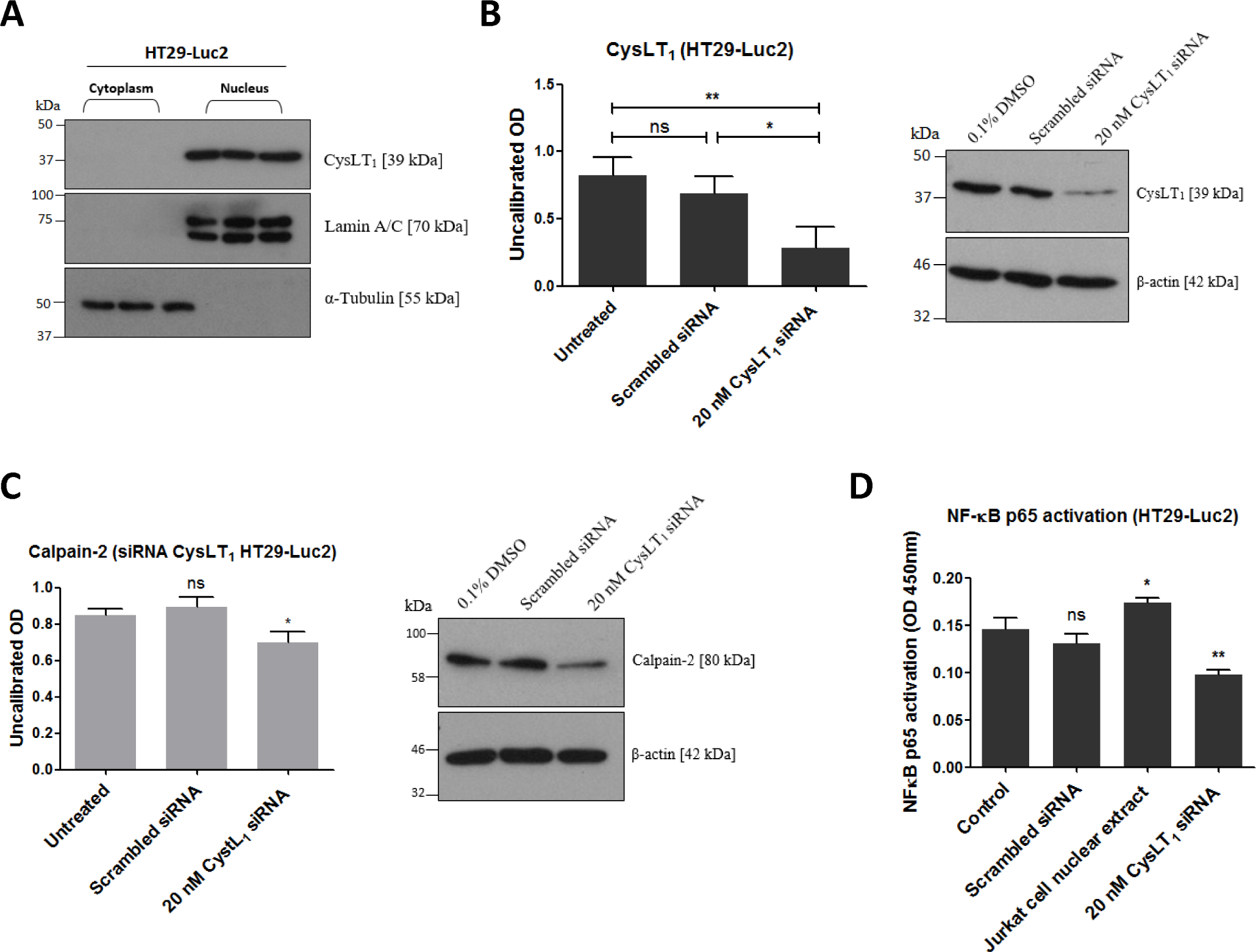
Effects of CysLT_1_ gene silencing in HT29-Luc2 cells. **A.** CysLT_1_ is expressed in the nucleus of HT29-Luc2 colorectal cancer cells. **B.** A unique 27mer siRNA efficiently silenced CysLT_1_ in HT29-Luc2 cells shown by reduced CysLT_1_ protein expression after 48 hours. **C.** Cell lysates isolated from HT29-Luc2 cells treated with 20 nM CysLT_1_ siRNA had reduced levels of calpain-2 protein and **D.** reduced levels of activated NF-ĸB p65. Statistical analysis was performed by student’s *t*-test. Error bars are mean +S.E. *, *p*<0.05; **, *p*<0.01; ns; not significant.

### Quininib analogues alter the secretions of pro-angiogenic factors from human ex vivo colorectal tumour explants

Human tumour explants representing Dukes’ stage A, B and C colorectal cancer were cultured in the presence of 10 μM Q8 for 72 hours. Multiplex ELISAs determined the levels of pro-angiogenic growth factors and inflammatory mediators secreted from the patient explants into tumour conditioned media (TCM) in n=15 patients. The angiogenic panel consisted of VEGF, VEGF-C, VEGF-D, PIGF, bFGF, FLT-1 and TIE-2 while the 10-plex inflammatory panel included IL1-þ, IL-2, IL-4, IL-6, IL-8, IL-10, IL-12p70, IL-13, IFN-γ and TNF-α. Q8 treatment of CRC *ex vivo* explants showed no significant difference in expression of VEGF, PIGF (**Figure 3**), VEGF-C, VEGF-D, and FLT1 (**Supp. Figure 2**) from the angiogenic panel. A significant reduction in TIE-2 expression was observed (*p*<0.01). Q8 treatment of CRC *ex viv*o explants showed no significant difference in expression of IL-4 (**Figure 3**), IL1-þ, IL-2, IL-6, IL-8, IL-10, IL-12p70, IL-13, IFN-γ and TNF-α (**Supp. Figure 2**) in the inflammatory panel. Analysis of the adhesion molecules ICAM-1 and VCAM-1 in n=17 patients showed a significant reduction in VCAM-1 (*p*<0.05) but not ICAM-1 expression following treatment with Q8 (**Figure 3**).

**Figure 3.**
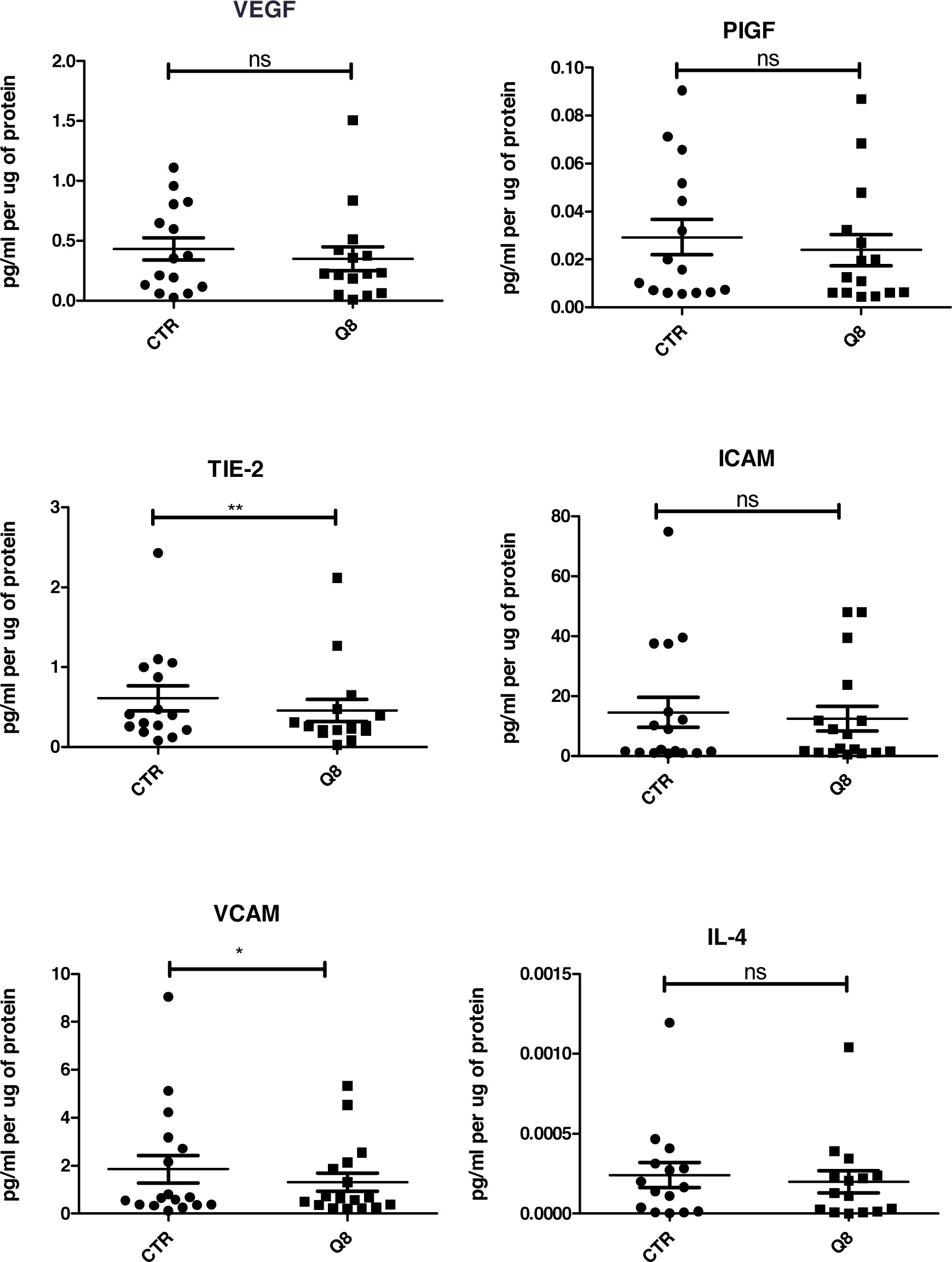
ELISA of tumour conditioned media from patient samples showing 72 hr treatment of 10 uM Q8 significantly reduces the expression of the angiogenic marker TIE-2 and the adhesion molecule VCAM compared to Control (0.1% DMSO). Statistical analysis was performed by Wilcoxon signed rank test. Q8 significantly reduced TIE-2 and VCAM expression compared to control (0.1% DMSO). *,p<0.05;**,p<0.01. VEGF, PIGF, ICAM and IL-4 were not significant with IL-4 showing a trend towards significance (p=0.0554). n=15 in all cases except VCAM and ICAM (n=17)

### Baseline expression levels correlate with patient nodal status

To determine if expression levels of the profiled angiogenic or inflammatory markers correlate with clinical patient parameters (n=14), spearman correlations were conducted using statistical software R (16) and graphical visualizations generated using the Corrplot package for R (**Figure 4A**) (17). A complete list of spearman correlation values can be found in **Supplementary Table 1A**. The baseline expression of PIGF, TIE-2, ICAM-1 and VCAM-1 secreted from the human colorectal explants correlated with patient nodal status (r = 0.60, 0.70, 0.62 and 0.67 respectively, **Figure 4C**). Patients with higher secretion of PIGF, TIE-2, ICAM-1 or VCAM-1 from their tumour explants displayed higher nodal status, reflective of a more aggressive cancer phenotype. Following the addition of Q8 (10 μM), the expression level of each analyte was compared to its baseline (control) expression level to determine the relative fold change. Relative Q8 fold change levels were again correlated to patient clinical parameters. In this instance the following associations were determined; FLT-1 correlations were greatly reduced with bFGF, PIGF, TIE-2, VEGF, IL-4, TNF-α, ICAM-1 and VCAM-1, while IL1-þ correlations were greatly reduced with IL-2, IL-6, IL-8 and IL-10 **Figure 4B, Supplementary Table 1B**).

**Figure 4.**
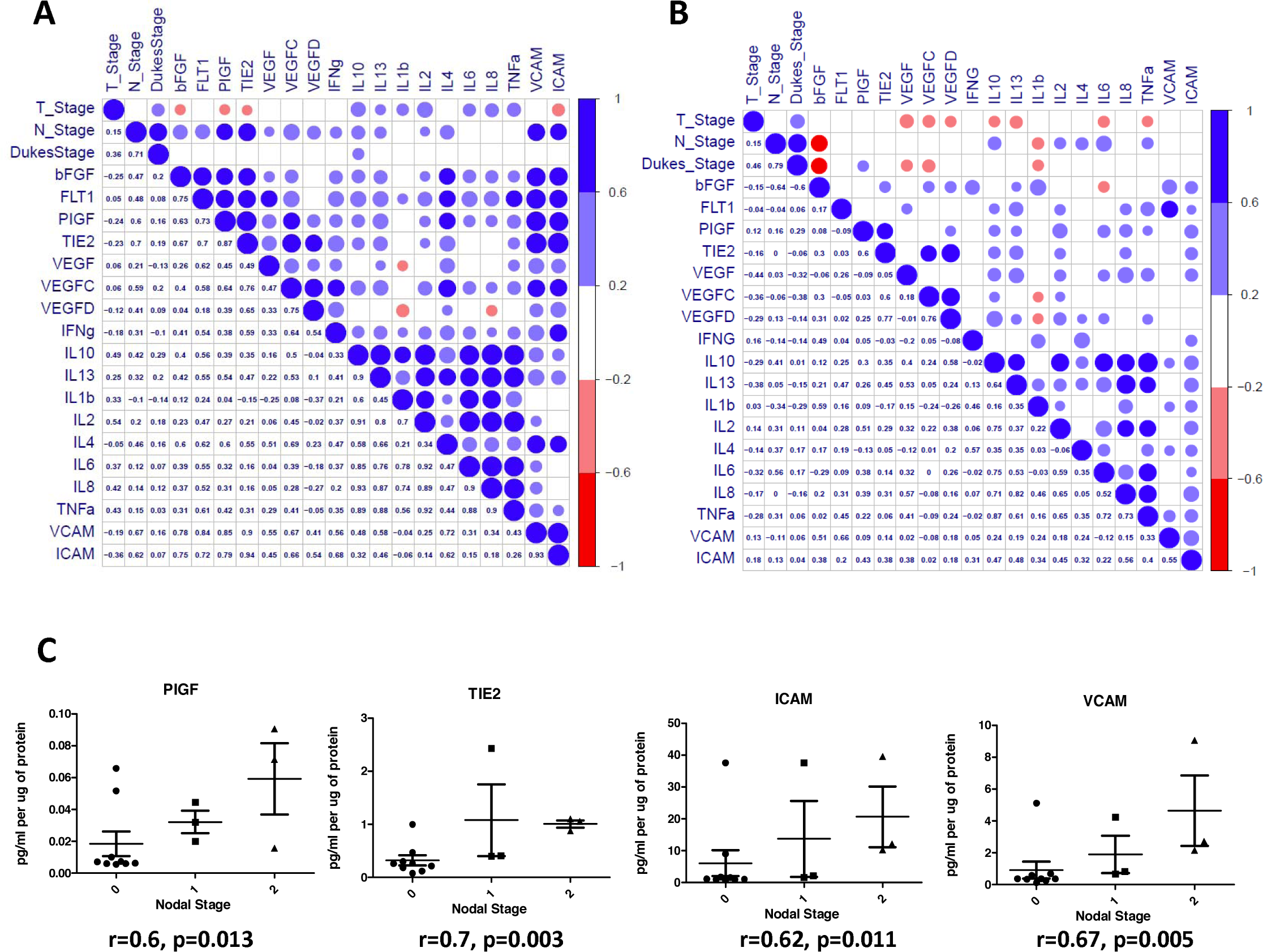
Correlation of angiogenic and inflammatory analytes and clinical parameters. Secreted baseline (control) levels of 18 ELISA analytes were compared in n=14 patients to determine if correlations existed between angiogenic and inflammatory analytes and between patients clinical parameters (Tumour Stage (T_Stage), Nodal Stage (N_stage) and Dukes Stage (A). Following the addition of Q8 (10uM), the value of each analyte was compared to its baseline (0.1% DMSO control) expression level to determine the relative fold change. Relative Q8 fold change levels of 18 ELISA analytes were compared in n=14 patients to determine if correlations existed between angiogenic and inflammatory analytes and between patients clinical parameters as in A, (B). The lower left triangle denotes spearman correlation values for each comparison while the upper right triangle represents correlation value. A blue circle denotes a positive correlation (>0.6), while a red circle denotes a negative correlation (<-0.6). (B) (C) Nodal stage was found to correlate with baseline levels of PIGF, TIE2, ICAM, and VCAM expression levels. A complete list of correlation values can be found in Supplementary Table 1.

### Q8 is safely tolerated in Balb/C mice

As Q8 has anti-angiogenic action *ex vivo*, and affects clonal cell survival *in vitro*, we evaluated its safety profile in advance of *in vivo* efficacy in CRC xenografts in collaboration with a pathologist (RD). During Q1 and Q8 treatment (6 intraperitoneal doses) no clinical adverse events were detected up to 50 mg/kg. At 100 mg/kg Q1 or Q8, significant weight loss occurred in ∼20% animals and piloerection in all animals. Hence, the maximum tolerated dose for Q1 and Q8 was determined to be 50 mg/kg (**Supp. Figure 1 A**). Histological examination of organ tissue from mice treated with 10% DMSO (vehicle) revealed that all specimens were histologically within normal limits (**Supp. Figure 1 B**). All tissue specimens in the 50 or 100 mg/kg Q1 group were noted as histologically within normal limits, however, vascular calcifications were identified in the kidneys (two animals at each dose) and in the heart (one animal at each dose) (**data not shown**) (**Supp. Figure 1 C**). All tissue specimens in the 50 or 100 mg/kg Q8 group were histologically within normal limits except for vascular calcification identified in the kidney of one animal and foci of non-specific, mild chronic lobular inflammation identified in the liver of one animal treated with 100 mg/kg Q8 **(data not shown)** (Supp. Figure 1 D). All tissue specimens were histologically within normal limits in the 25 mg/kg Q8 group (Supp. Figure 1 E). In summary, the MTD of Q1 was confirmed at 50 mg/kg and the MTD of Q8 was determined at 50 mg/kg. There were no significant histological differences between organs of the control group, Q8 (25, 50 and 100 mg/kg) and Q1 (50 and 100mg/kg) groups.

### Q8 reduces tumour volume and bioluminescence in HT29-Luc2 xenografted mice

To determine if Q8 significantly reduces tumour growth in vivo, and to compare to quininib (Q1), CRC-specific murine xenograft models were created by subcutaneous injection of HT29-Luc2 cells using immune-compromised mice (Balb/C nu/nu). Palpable tumours were evident and measurable with a digital calipers 7 days post subcutaneous implantation of 2.5 x 10^6^ HT29-Luc2 cells per mouse. 3 drug groups (each with 6 animals) were treated by intraperitoneal injection every 3 days and included; 25 mg/kg Q1, 25 mg/kg Q8 or vehicle (10% DMSO). Treatment with 25 mg/kg Q1 significantly decreased tumour volume (**Figure 5 A**), as measured using callipers, compared to vehicle control (average final measurements: 633.59 mm^3^ vs. 814.70 mm^3^, *p*<0.05). Treatment with 25 mg/kg Q8 also significantly reduced tumour volume (Figure **5 A & C**) compared to vehicle control (average final measurements: 553.31 mm^3^ vs 814.70 mm^3^, *p*<0.001). On the final day of the study, the bioluminescent activity of tumours from all animals was assessed following intraperitoneal luciferin injection. Bioluminescence of tumours was lowest in the Q8 group with Q8 bioluminescence being significantly lower compared to that of the control (average bioluminescence 1.24 x 1010 photons/sec vs. 2.33 x1010 photons/sec, *p<0.05*) on day 41 (**Figure 5 B & C**). There was no statistically significant difference in bioluminescence between control and Q1 treatment groups on day 41 of the study. There was no statistically significant difference in bioluminescence between Q8 and Q1 treatment groups on day 41 of the study. The *ex vivo* tumours, livers and lungs of all mice were imaged using the IVIS^®^ spectrum to search for evidence of HT29-Luc2 cell metastasis but no bioluminescence was observed in the lung or liver tissue taken from either control, Q1 or Q8 treated mice indicating that the HT29-Luc2 cells of the primary subcutaneous tumour had not metastasized (**Figure 5 D**). In summary, Q8 significantly reduced tumour volume and tumour bioluminescence in HT29-Luc2 xenografted mice compared to vehicle control.

**Figure 5.**
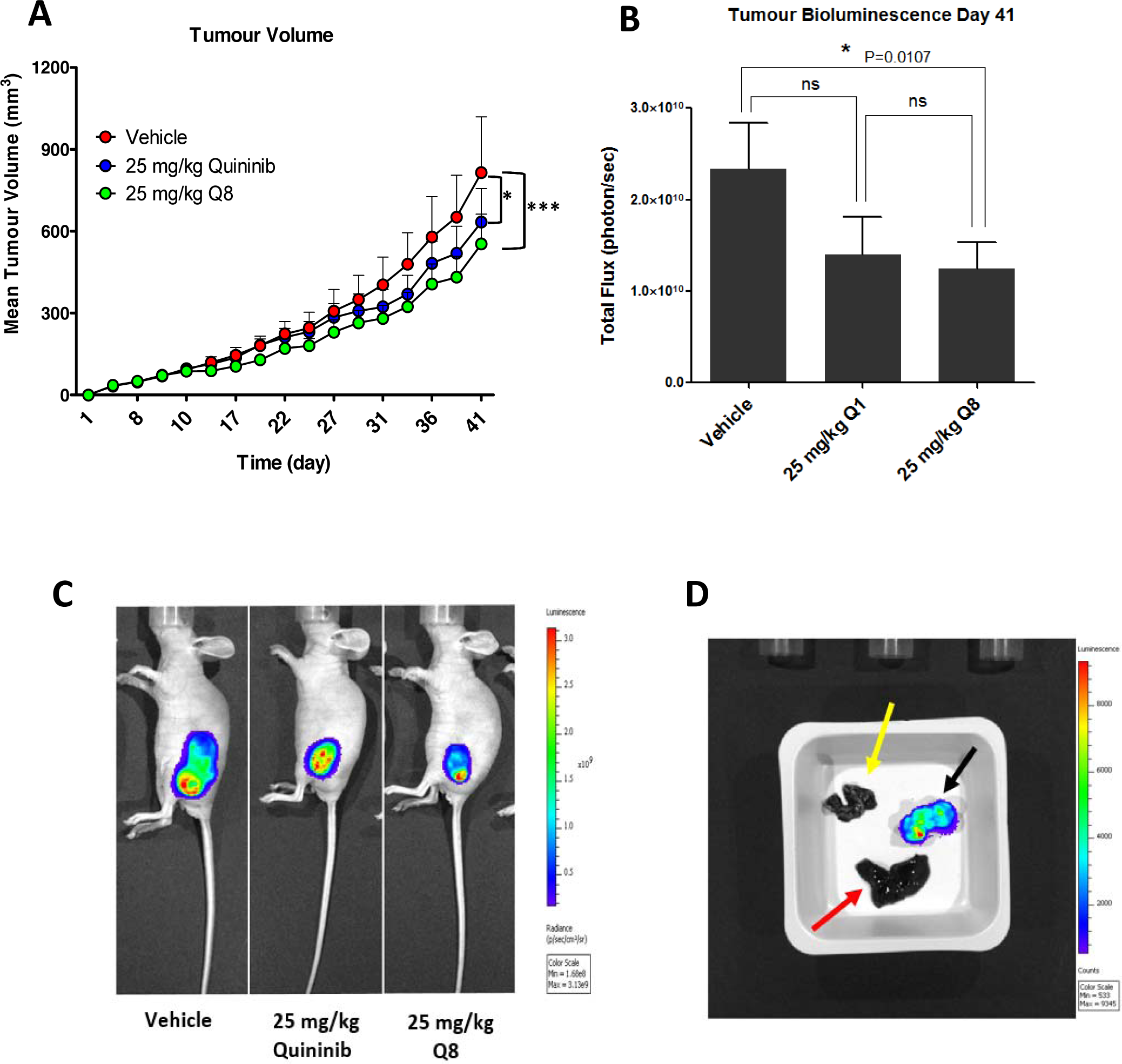
Q8 reduces tumour volume and bioluminescence in HT29-Luc2 xenografted mice. **A.** Effects of Q8 compared to vehicle control and Q1 on tumour volume in immune-compromised mice over 41 days, as measured using digital calipers (n=6 mice per group). **B.** Bioluminescent activity of tumours in xenografted mice on day 41 of the study, imaged using the IVIS^®^ spectrum. **C.** Images of mice during bioluminescent analysis of tumours from control, Q1 and Q8 treated mice. **D.** *Ex vivo* analysis of tumour (black arrow), liver (red arrow) and lungs (yellow arrow) from a control treated mouse for bioluminescent activity (metastasis of primary tumour cells). No metastasis was evident in the liver or lungs of all animals in the study at day 41. Statistical analysis was performed by ANOVA with Dunnett’s post hoc multiple comparison test. Error bars are mean +S.E. *, *p*<0.05; ***, *p*<0.001; ns; not significant.

### The effect of Q8 on the expression of angiogenic, apoptotic and proliferative markers in mouse xenograft tumour tissue

Immunohistochemical analysis of post-mortem xenograft samples revealed the level of proliferation positivity, Ki-67, the apoptotic marker caspase 3 and TIE-2 were not significantly decreased in the 25 mg/kg Q8 treatment group compared to vehicle control (all *p* values > 0.05, **Figure 6A-C**). However, following Q8 treatment, the levels of calpain were significantly decreased (**Figure 6D**, *p*<0.0001).

**Figure 6.**
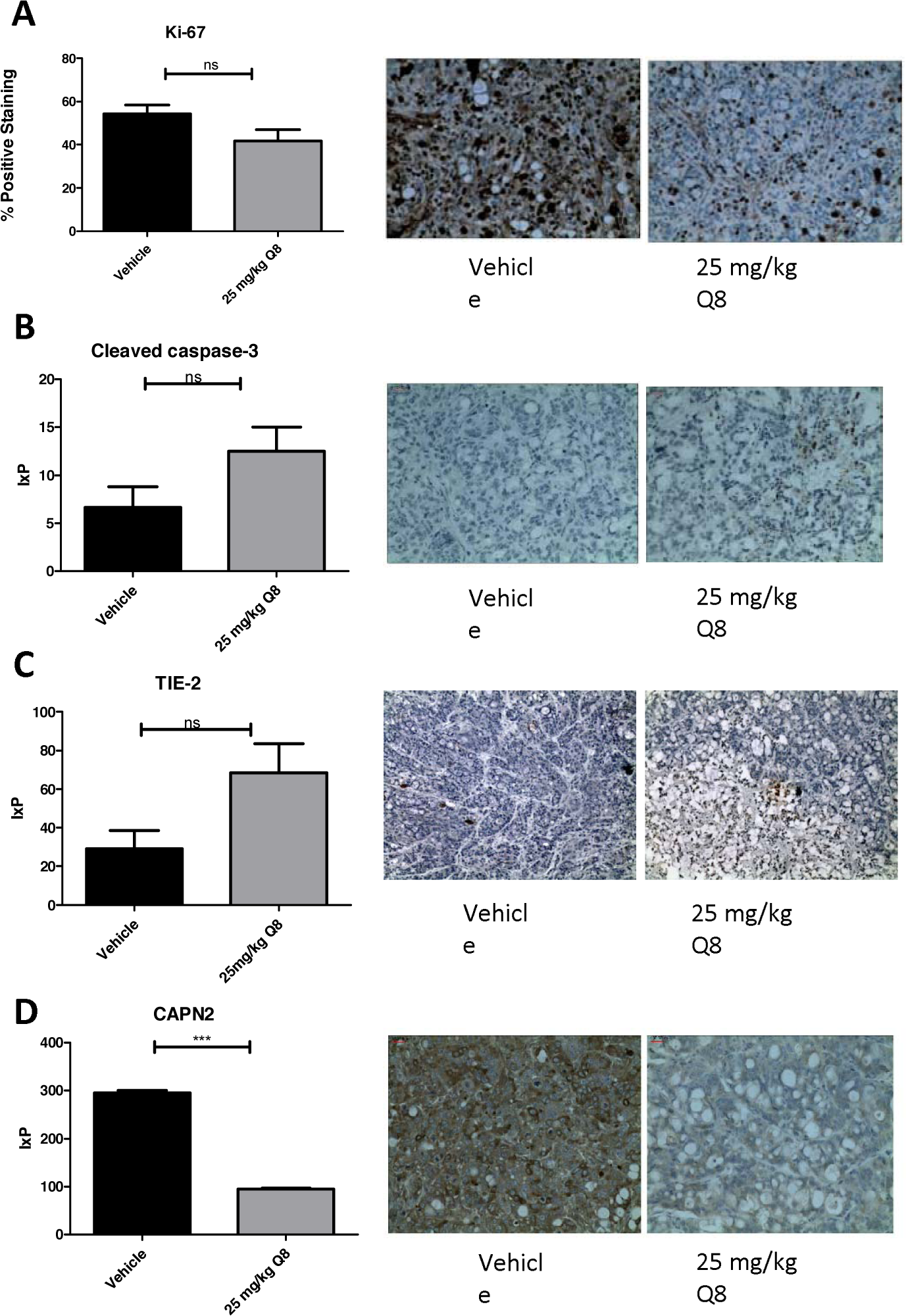
Immunohistochemistry analysis of Q8 in mouse xenograft tumour tissue. **A.** 25 mg/kg Q8 showed no significant decrease in expression of proliferation marker Ki-67 in xenograft tumour tissue compared to vehicle (p=0.0924). **B.** 25 mg/kg Q8 showed no significant difference in expression of apoptotic marker cleaved caspase-3 in xenograft tumour tissue compared to vehicle (p=0.1048). **C.** 25 mg/kg Q8 did not significantly increase TIE-2 expression compared to vehicle (p=0.0518). **D.** 25 mg/kg Q8 significantly reduces expression of calpain-2 in mouse tumour xenograft tissue compared to vehicle (p<0.001) Statistical analysis was performed using unpaired student’s *t*-test. Error bars are mean +S.E.); ns; not significant,***; p<0.001. IxP = Staining intensity * % Positive staining

## Discussion

Based on the robust anti-angiogenic activity of 1,4 dihydroxy quininib (Q8) (13) in comparison to the other analogues tested, it was the analogue chosen to further test across different models. This study demonstrated that the quininib analogue, 1-4 dihydroxy quininib (Q8), a CysLT_1_ antagonist showed significant anti-cancer properties in CRC based on *in vitro, ex vivo* and *in vivo* assessment of this drug. Overall, Q8 acts in an alternative pathway, non-redundant to the VEGF pathway, and may represent an alternative treatment strategy to counteract anti-VEGF resistance in CRC.

Our *in vitro* work has shown that Q8 alters CRC cell survival and that the CysLT_1_ nuclear expression in HT29-Luc2 cells regulates downstream effectors of calpain-2 and NF-ĸB. CysLT_1_ agonism promotes NF-ĸB p65 activity (13). NF-ĸB is a well-established enhancer of tumorigenesis, promoting epithelial-mesenchymal transition (31), cell proliferation and VEGF-induced tumour vascularization (18). 1,4-dihydroxy quininib (Q8) is an antagonist of CysLT_1_ and reduces the activation and DNA binding of NF-ĸB p65 in human endothelial cells (13). To corroborate these findings in CRC models, we analysed the expression and function of CysLT_1_ in HT-29-Luc CRC cells. CysLT_1_ is abundantly expressed in the nucleus of HT29-Luc2 cells and CysLT_1_ gene silencing inhibited NF-ĸB p65 activity in these CRC cells. Similarly, calpain-2 is a known regulator of VEGF mediated angiogenesis (19). Calpain-2, also known as m-calpain when bound to its regulatory subunit calpain-S1, is over expressed in colorectal tumour cells (20,21). The increased activity and expression of calpains in cancer cells during tumorigenesis (22) is largely attributed to growth factor signalling and the oncogenes v-Src (23) and c-Myc (24). Both Src and c-Myc proteins contribute to the malignant phenotype of HT29 cells (25-27). In endothelial cells 1,4-dihydroxy quininib reduces calpain-2 expression (13) and here this is corroborated in HT29-Luc2 CRC cells, as evidenced by decreased calpain-2 expression following CysLT_1_ gene silencing.

Human cell lines do not recapitulate the complexity of the tumour microenvironment. Human tumours cultured *ex vivo* better recapitulate the cellular heterogeneity associated with tumorigenesis and allow the assessment of drugs on protein secretions *ex vivo* (28). We analysed the anti-CRC effect of 1,4-dihydroxy quininib in human CRC patient explant tissue and assessed how Q8 affected the protein secretion levels of a panel of angiogenic and inflammatory proteins. Q8 specifically affected the secretions of soluble VCAM-1 and TIE-2 proteins and did not alter proteins linked to VEGF signalling. VCAM-1 is a membrane bound or circulating cytokine activated endothelial cell adhesion molecule which binds to leukocyte integrin VL-4 (very late antigen-4) on leukocytes, signalling their migration to inflammatory regions (29). VCAM-1 regulates tumour angiogenesis by enabling endothelial cell adhesion. In gastric tumours, VCAM-1-positive tumours are more invasive, later disease and more metastatic compared to VCAM-1 negative tumours (30). VCAM-1 acts downstream of the Angiopoietin-TIE pathway. Kim *et al.* demonstrated that in endothelial cells Angiopoietin-1 (ANGPT1) exerts its effect by receptor binding of TIE-2 and that ANGPT-1 suppressed the VEGF induced expression of adhesion molecules E-selectin, ICAM-1 and VCAM-1 (31). As previously stated, TIE-2 is not only expressed in endothelial cells but also in pericytes and TIE-2 expressing monocytes. In a subset of bone marrow fibroblasts, ablation of VCAM-1 in a mouse model indicated their developmental origin was from a TIE-2 expressing progenitor (32). Due to the significant reduction in TIE-2 and VCAM-1 expression we postulate that Q8 may be acting on the angiopoietin-TIE-2 signalling pathway (Figure 7).

**Figure 7.**
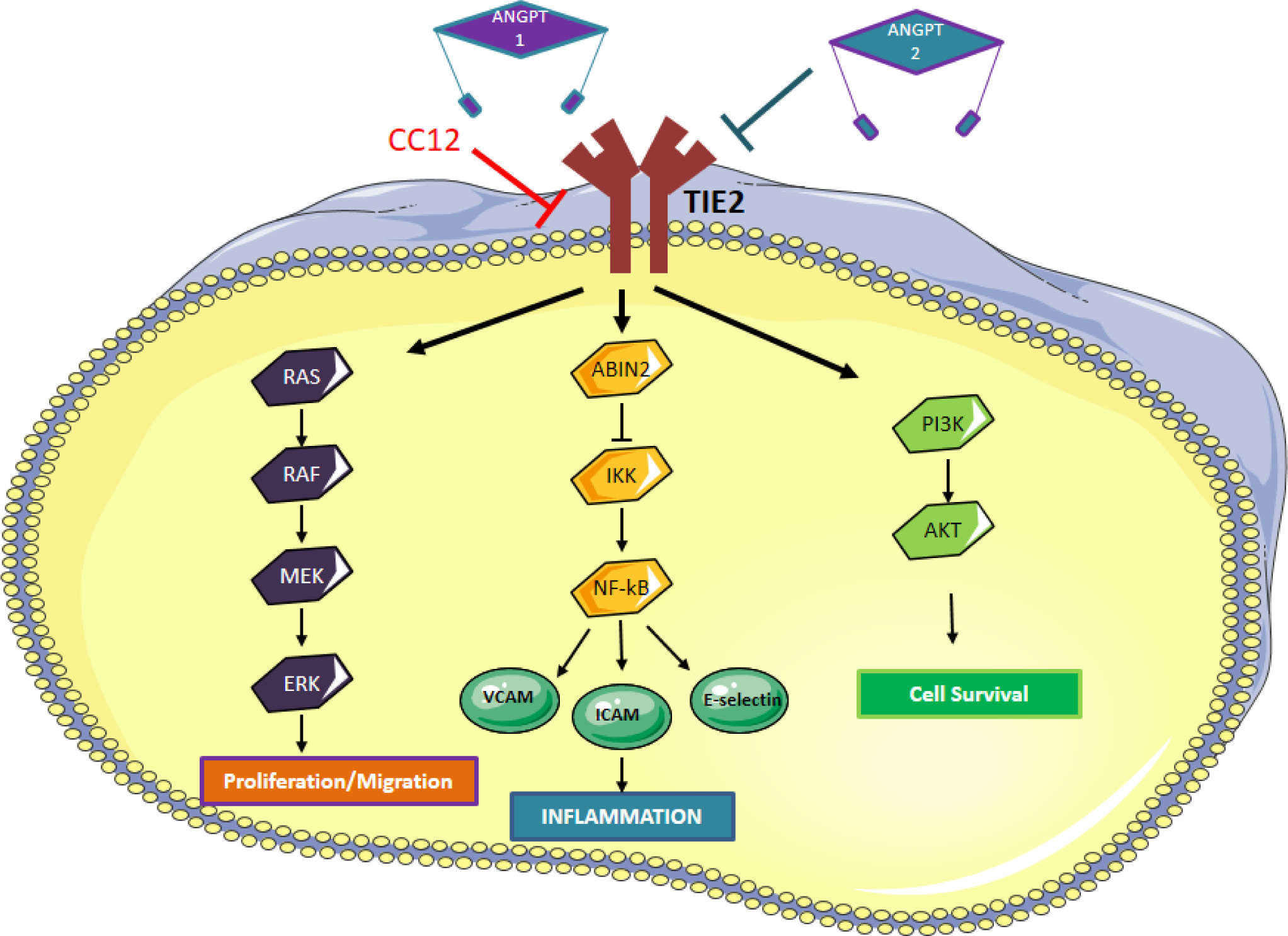
Angiopoietin-TIE signaling pathway. Binding of ANGPT1 to TIE2 induces autophosphorylation of TIE2 and the subsequent activation of multiple downstream pathways involved in migration, inflammation and cell survival. ANPT2 acts as an antagonist to ANGPT1 by competing with ANGPT1 to prevent phosphorylation of TIE2, however, ANGPT2 can induce TIE2 signaling in the absence of ANGPT1 or via overexpression of ANGPT2. We hypothesise that Q8 is acting on the Angiopoietin-TIE signaling pathway and downstream inflammatory network through its reduction in TIE2 and VCAM expression levels.

Following the *proof-of-concept* studies in CRC cells, *in vitro*, and patient tumour explants, *ex vivo*, we tested the hypothesis that 1,4-dihydroxy quininib could attenuate growth of CRC tumour xenografts *in vivo*. The MTD of 1,4-dihydroxy quininib was determined at 50 mg/kg and repeated administration every 3-4 days over 3 weeks resulted in no significant histological changes. Notably at 25 mg/kg, 1,4-dihydroxy quininib significantly reduced tumour volume and tumour bioluminescence compared to vehicle control. Furthermore, 1,4-dihydroxy quininib reduced the expression of the pro-angiogenic factor calpain-2, corroborating the earlier findings we detected in HT29-Luc2 cells. The reduction in calpain-2 expression in xenograft tumours may serve as a novel outcome to complement existing anti-VEGF therapies for colorectal cancer such as Bevacizumab. Calpains play a poignant role in tumour angiogenesis by regulating the survival and migration of endothelial cells (20,33). Inhibition of calpains represents a novel anti-angiogenic strategy in cancer but such inhibition needs to be highly specific as m-calpain enables endothelial cell migration but µ-calpain enables endothelial cell apoptosis and vascular regression (34). VEGF stimulates endothelial cell migration by enhancing m-calpain (calpain-2) expression and activity allowing tail retraction and this process can be reversed using calpastatin, an inhibitor of calpain-2 (19). Although calpain-2 (m-calpain) orchestrates neovascularization, µ-calpain is responsible for cell apoptosis and angiogenic regression (34), therefore, select and endothelial specific inhibition of m-calpain would be of great importance in a cancer setting. Calpain-2 also plays a role in tumour and stromal cell invasion during disassembly of focal adhesions at the rear of the tumour or stromal cell (35). While immunohistochemical analysis of human colorectal adenocarcinomas have previously shown strong expression levels of TIE-1 and TIE-2, which correlated with several clinicopathological factors (36) we observed weak staining in our xenograft tissue and no change in expression following Q8 treatment *in vivo*. This is in contrast to the changes we detect in human *ex vivo* explants treated with Q8. In the human model the functional soluble TIE-2 was significantly reduced with Q8. The levels of secreted TIE-2 and the level of total TIE-2 expression in tissue are measuring TIE-2 at different levels but the secreted TIE-2 is suggested to be more functional.

Drug resistance remains a major problem with current therapies such as bevacizumab, sunitinib and sorafenib, which target the VEGFR system, the chief regulator of angiogenesis. Numerous preclinical studies of angiopoietin-TIE targeted treatment in oncology and ocular vascular diseases suggest complementation with VEGF targeted therapy (6). Many studies aim to block the binding of ANGPT1/ANGPT2 to TIE-2, thereby, inhibiting tumour growth. TIE-2 functionality and activation or indirect inhibition of its expression is context-dependent. Inhibition of ANGPT2 and the subsequent activation of TIE-2 by the ANG2 Binding TIE-2 Activating Antibody (ABTAA) improves vascular stabilization and drug delivery in mouse models of orthotopic glioma, subcutaneous Lewis lung carcinoma and spontaneous mammary cancer (37). TIE-2 expressing macrophages induce vessel destabilization, promote intravasation of tumour cells in the primary tumour (6) and would consequently warrant a reduction in TIE-2 expression. However, in pericytes, a pro-angiogenic effect was observed leading to enhanced tumour growth following the deletion of TIE-2 (7). The importance of the ANGPT-TIE-2 signalling pathway which acts as a link between angiogenesis and inflammation positions it as an excellent target to modify therapeutically.

In summary, 1,4-dihydroxy quininib may be an alternative therapeutic approach for patients who become resistant to anti-VEGF drugs, possibly in combination with standard treatment for these patients.

## Experimental Procedures

### Cell culture

HT29-Luc2 Bioware^®^ Ultra human colorectal cancer cells were purchased from (PerkinElmer/cat: 124353) and were maintained at 37 °C / 5% CO_2_ in McCoy’s 5A medium (Gibco) with L-glutamine (Gibco) supplemented with 10% fetal bovine serum (FBS) (Gibco).

### MTT assay

3-(4,5-dimethylthiazol-2-yl)-2,5-diphenyltetrazolium bromide) dye was used to determine the cytotoxic effect of quininib and analogues on HT29-Luc2 cells. Cells were trypsinised using 2 ml TrypLE™ Express (Gibco) and centrifuged at 1,200 rpm for 4 mins at RT. Cell pellets were re-suspended in full medium and cells were counted and seeded into 96-well plates at a density of 10,000 cells/well. Cells were left to adhere for 24 hours and serum starved for 24 hours. Medium was removed from HT29-Luc2 cells and replaced with 10 μM drug solution, where 5-FU was used as a positive control and DMSO as a vehicle control. HT29-Luc2 cells were incubated for 24, 48, 72 and 96 hours with drugs. Drug solution was removed and wells were washed with PBS before adding 10 µl MTT (3-(4,5-dimethylthiazol-2-yl)-2,5-diphenyltetrazolium bromide) dye to cells for 4 hours at 37 °C. 100 µl 100% DMSO was added to wells to dissolve the formazan crystals which formed. The absorbance values were read at 570 nm using a SpectraMax^®^ M2 microplate reader.

### Colony formation assay

HT29-Luc2 CRC cells were maintained in McCoy’s 5A (Gibco) medium supplemented with 10% FBS (Gibco) at 37 °C/5% CO_2_. Cells were washed in DPBS and trypsinised using 2 ml TrypLE™ Express (Gibco). Cells were centrifuged following detachment at 1,200 rpm for 4 minutes. Cell pellets were re-suspended in full medium and a cell count was performed using a haemocytometer. 1.5 x10^3^ cells were seeded per well of a 6-well plate and left to adhere for 24 hours. Cells were then treated with 10 μM or 20 μM of quininib (Q1), Q8, Q18, Q22, P4 or P18 for 24, 48, 72 and 96 hours. The drugs were removed and cells were grown in fresh media for 10 days in total, clones were then fixed using 4% paraformaldehyde and stained by incubating with 0.5% crystal violet solution (Pro-Lab diagnostics PL7000) at RT for 2 hours. Clone counting was performed using the ColCount™ system (Oxford Optronix).

### CysLT_1_ expression

HT29-Luc2 cells were seeded at 250,000 cells per well of a 6-well plate and following adherence, cells were serum starved for 24 hours. To assess CysLT_1_ expression, cytosolic and nuclear proteins were extracted from cells. Cells were washed in ice-cold PBS and 200 µl Buffer A (10 mM HEPES pH 8, 1.5 mM MgCl_2_, 10 mM KCl, 200 mM sucrose, 0.5 mM DTT, 0.25% IGEPAL (octylphenoxypolyethoxyethanol) and 1x protease inhibitor) was added to each well and left for 5 minutes. Cells were scraped into an eppendorf and were left to swell on ice for 15 minutes. 12.5 ul of 5% NP-40 (IGEPAL) (Sigma) was added to each 200 ul sample to give a final concentration of 0.3% NP-40) and samples were vortexed for 10 seconds each. Cell lysates were centrifuged at 12,000 rpm for 2 minutes at 4 °C. Supernatant was removed from each sample and stored at −20 °C as cytoplasmic fractions. The remaining pellet was washed in ice-cold PBS to remove any remaining traces of cytoplasmic fraction and 50 ul of Buffer B (20 mM HEPES pH 8, 420 mM NaCl, 0.2 mM EDTA, 1.5 mM MgCl_2_, 0.5 mM DTT, 25% glycerol and 1x protease inhibitor) was added to the remaining pellet and the sample was sonicated for 5 seconds. Each sample was centrifuged at 14,000rpm for 10 min at 4 °C, and the supernatant was stored as nuclear fraction at −80 °C. Following treatment of HMEC-1 cells with 10 µM Q1, Q8 or montelukast for 5 hours, total protein was extracted from cells using ice-cold RIPA lysis buffer supplemented with sodium fluoride (Sigma), β-glycerol-phosphate (Sigma) protease inhibitor (Sigma) and phosphatase inhibitor (Sigma). Cells were washed with ice-cold PBS and 100 µl ice-cold RIPA lysis buffer was added to each plate well. Cells were scraped into eppendorfs and left at 4 °C on a roller for 30 minutes to completely lyse the cells. Samples were centrifuged at 12,000 rpm for 15 minutes at 4 °C and supernatants were stored at −80 °C as whole cell lysates. Protein concentrations were determined using the BCA protein assay kit (Fisher) or Bradford assay reagent (Fisher) (for cytoplasmic/nuclear proteins). Expression of proteins of interest were determined by immunoblotting. 20 µg of protein was prepared in 5X sample buffer and separated by 12% SDS/PAGE, transferred to PVDF membranes (Millipore), and probed with primary antibodies (CysLT_1_: Abcam, 1:2000, Lamin A/C: Cell Signaling 1:1000, Alpha tubulin: Santa Cruz 1:200, calpain-2: Aviva Systems Biology.

### siRNA knockdown of cysteinyl leukotriene receptor 1 gene in HT29-Luc2 cells and downstream calpain-2 and NF-ĸB protein analysis

To investigate the downstream effects of silencing CysLT_1_, HT29-Luc2 cells were seeded in 6-well plates and grown overnight in full medium. When the cells reached ∼65% confluency the next day, a unique 27mer siRNA duplex targeting CysLT1 transcripts (OriGene) or Trilencer-27 universal scrambled negative control (Origene) were added using SiTran1.0 transfection reagent (Origene) at a final concentration of 20 nM, as per the manufacturer’s protocol. After 6 hours, the cells were washed in PBS and medium was replenished. After 48 hours, the cells were collected and processed for protein analyses by immunoblotting. The unique 27mer siRNA 3’-GCAUUGAUCACCAUAAGAAGGAACA-5’ efficiently knocked down CysLT_1_ transcripts. Total protein was extracted from cells using RIPA lysis buffer supplemented with sodium fluoride (Sigma), β-glycerol-phosphate (Sigma) protease inhibitor (Sigma) and phosphatase inhibitor (Sigma). Protein concentrations were determined using the BCA protein assay kit (Fisher). 20 µg of protein was prepared in 5X sample buffer and separated by 12% SDS/PAGE, transferred to PVDF membranes (Millipore), and probed with primary antibodies (CysLT_1_: Abcam, 1:2000, calpain-2: Aviva Systems Biology, 1:2000 or β-actin: Sigma 1:5000). Membranes were subsequently probed with anti-rabbit horseradish peroxidase (HRP)-labelled secondary antibody: (Cell Signaling, 1:2000, or anti-mouse horseradish peroxidase (HRP)-labelled secondary antibody: Cell Signaling, 1:2000). Protein bands were visualized by chemiluminescent detection using Pierce ECL western blotting substrate (Fisher). Optical densitometry was used to quantitatively measure levels of expressed protein using ImageJ software. To determine the effects of silencing the CysLT_1_ gene on levels of activated NF-ĸB p65, 20 µg of HT29-Luc2 cells lysates were also analyzed for levels of activated NF-ĸB p65 using a TransAM NF-ĸB p65 kit (Active Motif) as per the manufacturer’s instructions.

### Maximum tolerated dose study (MTD)

Determination of the maximum tolerated dose of Q1 and Q8 was carried out by injecting balb/C normal female mice intraperitoneally 25, 50 or 100 mg/kg of Q8 and 50 mg/kg or 100 mg/kg Q1 every 3 days for a maximum of 10 doses. Mice were administered vehicle (10% DMSO), Q1 (50 & 100 mg/kg) or Q8 (25, 50 & 100 mg/kg) intraperitoneally every 3 days for 28 days (if tolerated) to determine the maximum tolerated dose (MTD). Each treatment group consisted of 3 mice (n=3). Any score of 4 from multiple observations or a score of 3 from a single observation resulted in immediate euthanasia of the mouse by CO_2_ asphyxiation. As the 100 mg/kg dose was deemed toxic, the animals of concern were subsequently euthanized 3 days after the initial 100 mg/kg Q1 dose and 4 days after the initial 100 mg/kg Q8 dose according to ethical guidelines. All animals were euthanized at the end of the study and necropsy was performed. The following organs were removed from each mouse and stored in 10% neutral buffered formalin (Sigma): brain, heart, lungs, kidneys, liver, small intestine and large intestine. Organs were processed using a Tissue-Tek^®^ Tissue Processor and paraffin embedded. Tissue sections were cut, deparaffinized and stained with hematoxylin and eosin. Tissue sections were analysed for gross morphological abnormalities by our collaborating pathologist (RD).

### Q8 tumour xenograft study

To determine if Q8 has enhanced effects on reduction of tumour growth compared to Q1, (previously shown to reduce tumour growth in mice), both drugs were tested on tumour growth by creating an optimized CRC-specific murine xenograft model by subcutaneous injection of 2,500,000 HT29-Luc2 cells in immunocompromised mice (Balb/C *nu/nu*). HT29-Luc2 cells were grown to 80% confluence, were recorded at passage 11 (p11) before use in the *in vivo* Q8 study and were kept on ice before subcutaneous injections. The Q8 study consisted of 6 mice per treatment group (n=6). Q8 was administered to mice by intraperitoneal injection every three days. Palpable tumours were evident and measurable with digital calipers (Scienceware^®^ Digi-Max^™^ slide caliper/cat: Z503576) 7 days post subcutaneous implantation of the HT29-Luc2 cells. When palpable tumours measured ≥100 mm^3^, mice were injected with drugs intraperitoneally using a 26-gauge needle (26G) every three days for ∼5 weeks. A tumour volume sheet was used to record tumour measurements every second day. Tumour volumes were calculated as per the following formula: 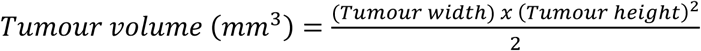. The appearance, weight, behaviour, respiration and level of tumour induced skin lesions of all mice were recorded every three days using the welfare score sheet (supplementary data). In keeping with ethical obligations outlined in AREC-P-11-42, if a tumour became necrotic, grew beyond 15 mm in diameter or became ulcerated, the animal was euthanized via CO_2_ asphyxiation. On day 41 of the study, one mouse receiving 10% DMSO was euthanized due to grade 2 wet ulceration of the subcutaneous tumour. Prior to euthanasia of this control mouse, all animals in the study were intraperitoneally injected with 150 mg/kg (150 μl) of D-Luciferin (15 mg/ml) (Perkin-Elmer) 10 minutes before being imaged using the IVIS^®^ Spectrum *in vivo* imaging system (Perkin-Elmer) to compare the bioluminescence of tumours between treatment groups. Following imaging, all animals were euthanized by CO_2_ asphyxiation and tumours were harvested by excision dissection and stored in 10% formalin or snapped frozen in liquid nitrogen for downstream histological analysis or protein extraction. To determine if metastases from subcutaneous tumours in mice had occurred, the liver and lungs were from removed from each animal of the Q8 tumour xenograft study immediately following euthanasia. This was done by making an incision through the peritoneum to expose the abdomen using a scissors. The liver and lung were gently cut out of the abdomen using a forceps and scissors. The organs were bathed in a 15 mg/ml solution of D-Luciferin and placed in a transparent petri-dish. The organs were placed beside the corresponding excised xenografted tumour tissue and imaged using the IVIS^®^ Spectrum *in vivo* imaging system.

### Immunohistochemistry

Xenograft slide sections were deparaffinized and rehydrated and tissue antigens unmasked using 1X Trilogy (Cell Marque) for 10 minutes in a pressure cooker. Sections were placed in fresh Trilogy solution for 5 minutes and in distilled water for a further 5 minutes. Slides were then placed in a 3% hydrogen peroxide solution (Sigma) for 30 minutes to quench endogenous peroxide activity. Slides were washed for 5 minutes in PBS three times and blocked to prevent non-specific antibody binding using goat serum (for rabbit antibodies) and horse serum (for mouse antibodies) (Vector Vectastain ABC mouse kit Vector Vectastain ABC rabbit kit). Primary antibodies were incubated with slide sections for 1 hour at RT at the following concentrations; Calpain-2 (Aviva Systems Biology) 10 µg/µl; Cleaved caspase 3 (Cell Signaling 1:50); Ki-67 (Dako 1:75); TIE-2 (Abcam, 1:100). Slides were washed for 5 minutes in PBS three times and incubated with biotinylated anti-rabbit or anti-mouse secondary antibodies (Vector Vectastain ABC kits) for 30 minutes at RT. Slides were washed for 5 minutes in PBS three times and incubated with ABC reagent for 30 minutes before staining with DAB (3, 3’-Diaminobenzidine tetrahydrochloride) (Sigma) and counterstaining nuclei with haematoxylin (Sigma). Slide sections were gradually dehydrated by incubating in 50%, 75%, 96% and 100% ethanol solutions, each for 3 minutes and cleared in Xylene (Sigma) overnight. Tissue sections were manually mounted with coverslips using DPX mounting medium (Sigma) and slides were left to dry overnight at RT. Slide sections were imaged at 20X magnification using a Nikon Eclipse E200 with OptixCam Summit series camera and OCView 7 software. The intensity of staining was graded semi-quantitatively in a blinded fashion by two observers (CB & AB) as negative “0”, weak “1”, moderate “2” or strong “3” and the positivity of staining was documented as 0% (no cells stained), 10% (some cells are stained but not 1/4 of cells stained), 25% (1/4 of cells stained), 50% (1/2 of cells stained), 75% (3/4 of cells stained), 90% (greater than 3/4 of cells stained but not all cells stained) or 100% (all cells stained). Immuno-histochemical staining for Ki-67 was performed in Beaumont hospital whereby deparaffinization, antigen retrieval and IHC staining for Ki-67 was performed on an automated platform (Bond™ III system – Leica MicroSystems™, Newcastle, U.K.). Slides stained with Ki-67 were scored for positivity only.

### Patient colorectal tumour explant ELISAs

Ethical approval for the use of human colorectal cancer explant tissue during experiments was granted from St James’s Hospital and Adelaide, Meath and National Children’s Hospital Institutional Review Board. Written informed consent was taken from all patients. Following the removal of a piece of colorectal tumour during patient surgery, part of the tissue was processed and paraffin embedded for histopathology analyses and the rest of the tissue was taken for explant culture experiments. The tissue was stored in a tissue wash buffer (PBS supplemented with 4 μg/ml fungizone (Gibco), 1% penicillin-streptomycin (Gibco) and 30 μg/ml gentamicin (Sigma). The explant pieces were thoroughly washed with additional tissue wash buffer within 30 minutes of the tissue being removed during surgery and stored in a 20% DMSO/HBSS buffer solution supplemented with 10% FBS, 2 μg/ml fungizone (Gibco) and 1% penicillin-streptomycin (Gibco)) and stored at –80 °C.. Each piece of human CRC tumour explant was thawed at RT and placed in a petri dish containing wash buffer (PBS supplemented with 4 μg/ml fungizone (Gibco), 1% penicillin-streptomycin (Gibco) and 30 μg/ml gentamicin (Sigma). The tissue was washed thoroughly and cut into smaller pieces. Smaller pieces of tumour were washed 3 times for 5 minutes in fresh wash buffer before being placed into wells of a 24-well plate containing Q8, diluted to 10 µM or 0.1% DMSO in warm RPMI-1640 medium supplemented with 10% FBS, 4 μg/ml fungizone (Gibco), 1% penicillin-streptomycin (Gibco) and 30 μg/ml gentamicin (Sigma). Explants were incubated for 72 hours at 37 °C and 5% CO_2_. Plates were wrapped in parafilm to prevent evaporation of drug-medium during the incubation period. After 72 hours, the remaining medium (TCM/tumour conditioning media) was stored at –20 °C and the residual explant tissue was snap-frozen in liquid nitrogen and stored at –80 °C. The levels of secreted angiogenic factors from human CRC explants was determined by carrying out ELISAs on the tumour conditioned media collected following explant culture. The human Angiogenesis Panel 1 containing VEGF, VEGF-C, VEGF-D TIE-2, FLT-1, PIGF Gen B, bFGF and Proinflammatory Panel 1 containing IFN-γ, IL1-þ, IL-2, IL-4, IL-6, IL-8, IL-10, IL12-p70, IL-13 and TNF-α Multiplex ELISA kits (Meso-Scale Discovery) were used to according to the manufacturer’s instructions. Levels of sVCAM-1 and sICAM-1 were detected by DuoSet ELISA (R&D Systems) as per manufacturer’s instructions. Patient tumour samples were homogenised in ice-cold T-PER lysis reagent (Fisher) (proprietary detergent with 25 mM bicine and 150mM sodium chloride (pH 7.6)) supplemented with 10 μl/ml protease inhibitor (Roche) and the BCA (Fisher) assay kit was used to quantify total protein extracted from explant tissue in μg/ml as per the manufactures instructions. 20 µg protein was analysed per sample using the calpain-2 ELISA according to the manufacturer’s instructions. Total protein was extracted from each piece of tumour explant tissue using ice-cold T-PER lysis reagent (Fisher) (proprietary detergent with 25 mM bicine and 150 mM sodium chloride (pH 7.6)) supplemented with 10 μl/ml protease inhibitor (Roche) to normalise angiogenic factor secretion from the ELISA to the amount of protein in each piece of tumour. Each piece of tissue was placed in a tube with 300 μl ice cold T-PER reagent and a 3 mm stainless steel bead (Qiagen). The tubes were placed in a TissueLyser II (Qiagen) for 2.5 minutes to completely homogenise the tissue, the lysate was centrifuged at 13,000 rpm at 4 °C for 5 minutes and the supernatant was used immediately or stored at − 80 °C. The BCA (Fisher) assay kit was used to quantify total protein extracted from explant tissue in μg/ml as per the manufacturer’s instructions.

### Statistical Analysis

Statistical analysis was carried out using Graphpad Prism 5 software. Corrplots were generated using the corrplot package (17) in R (16). Correlation of ELISA data with clinical parameters was conducted on n=14 patients as one patient’s IL-8 expression was below the level of detection. Therefore this patient was removed from all correlation analysis.

## Author Contributions

CTB, JOS and BNK designed the experiments. CTB, SK, AB, RD and EC conducted the experiments and analysed the data. CTB wrote the initial manuscript draft. SK, WMG, JOS and BNK interpreted the data and edited the manuscript to final version.

## Acknowledgments

The authors thank all patients who consented for their specimens and data to be used in this research. Figure 7 was prepared using Servier Medical Art PowerPoint image bank https://smart.servier.com/>

## Supplementary Info

**Supp. Figure 1.**
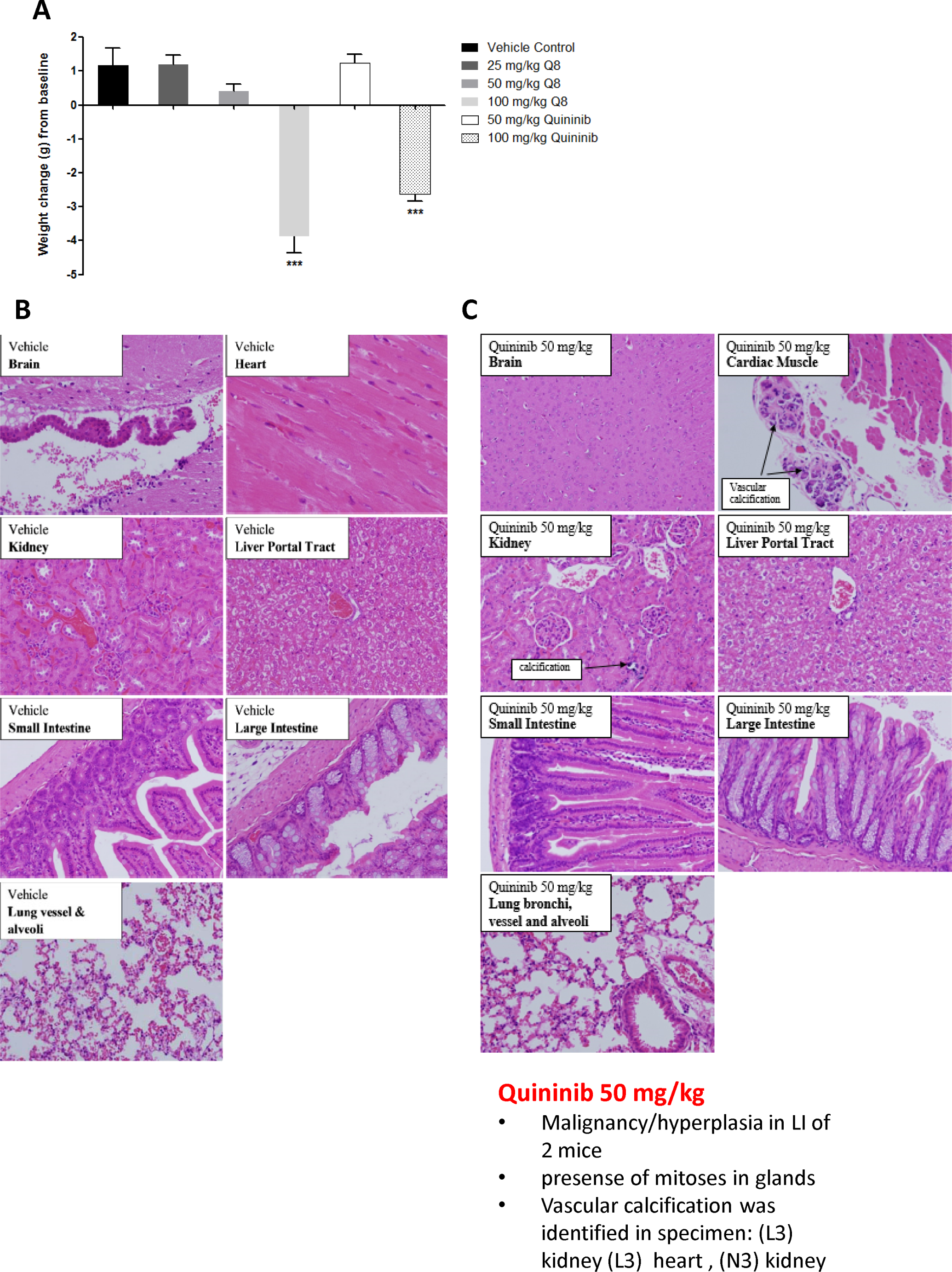
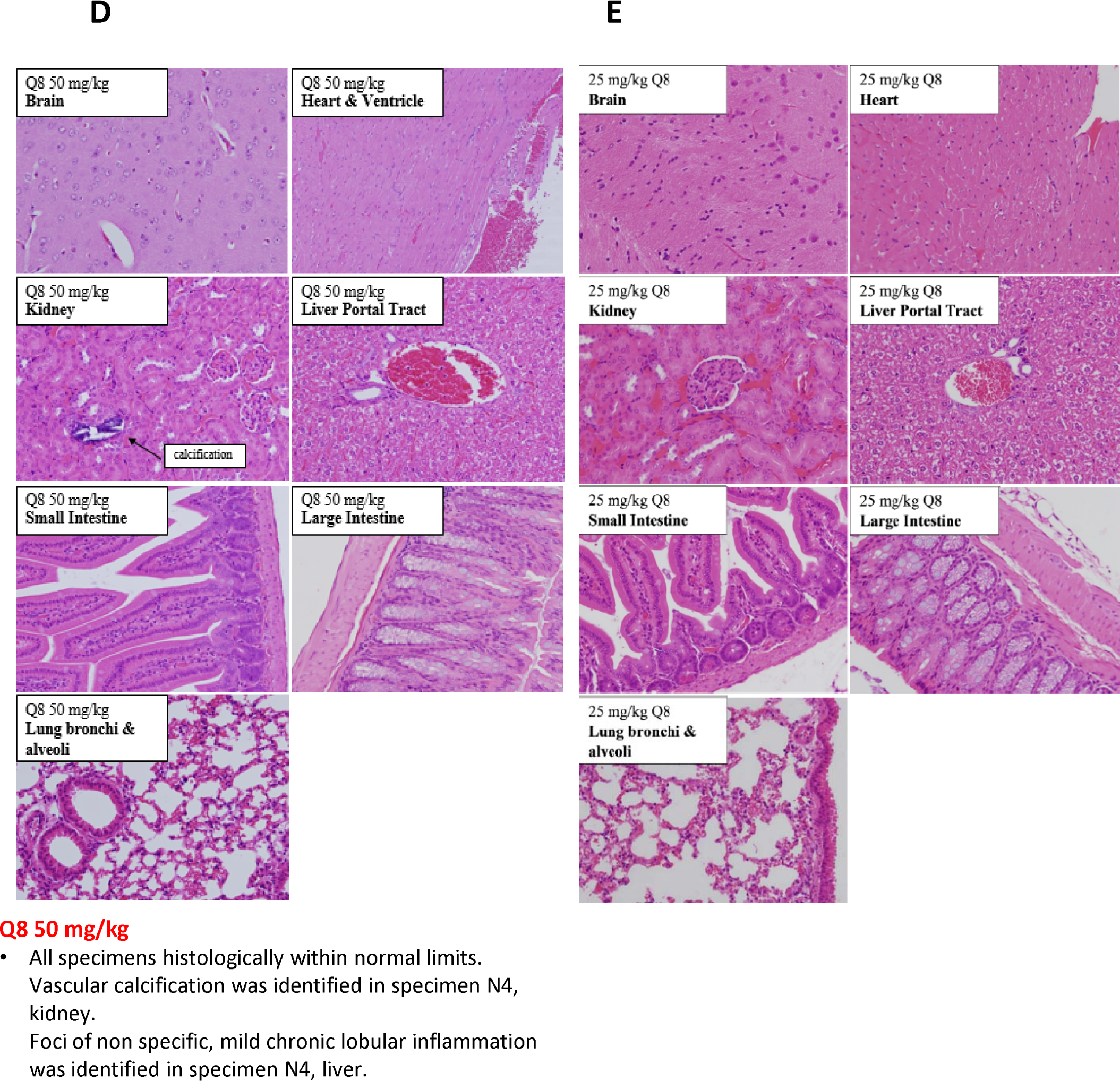
Weight changes of Balb/C wild type mice following i/p injection of Q1 and Q8 and histopathological analysis of organs obtained after treatment of mice with vehicle (10% DMSO) or 25 mg/kg Q8 showing absence of microscopic abnormalities. (A) Mice were treated with escalating doses of Q1 or Q8 (25 mg/kg – 100 mg/kg) every 3 days for 21 days. Drug administration ceased immediately and animals were euthanised if ≥20% body weight was lost or signs of discomfort such as piloerection, decreased respiration and vocalisation became apparent. Representative images of brain, heart, kidney, liver, small intestine, large intestine (colon) and lungs from Balb/C wild type mice treated with (B) vehicle control, (C). 50mg/kg Q1, (D) 50mg/kg Q8 and (E) 25 mg/kg Q8. All tissues were histologically within normal limits. Tissues were stained with hematoxylin & eosin. Magnification 20X. Statistical analysis was performed by ANOVA with Dunnett’s post hoc multiple comparison test. Error bars are mean +S.E. ***, *p*<0.001

**Supp. Figure 2:**
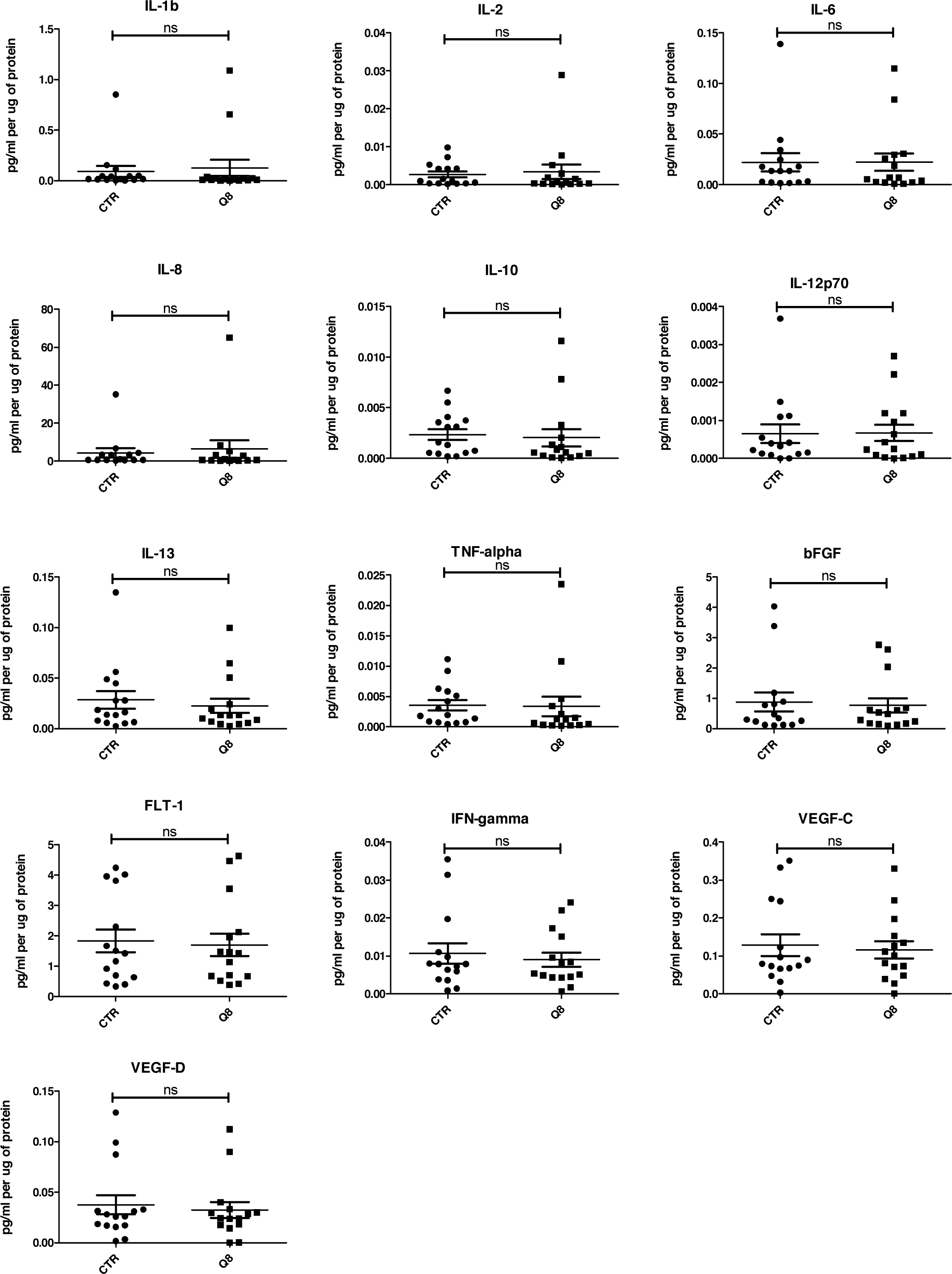
ELISA analysis of tumour conditioned media from resected patient colorectal tissue samples (n=15) treated for 72 hrs with 10 uM Q8 or 0.1% DMSO (CTR).

**Supplementary Table 1:**
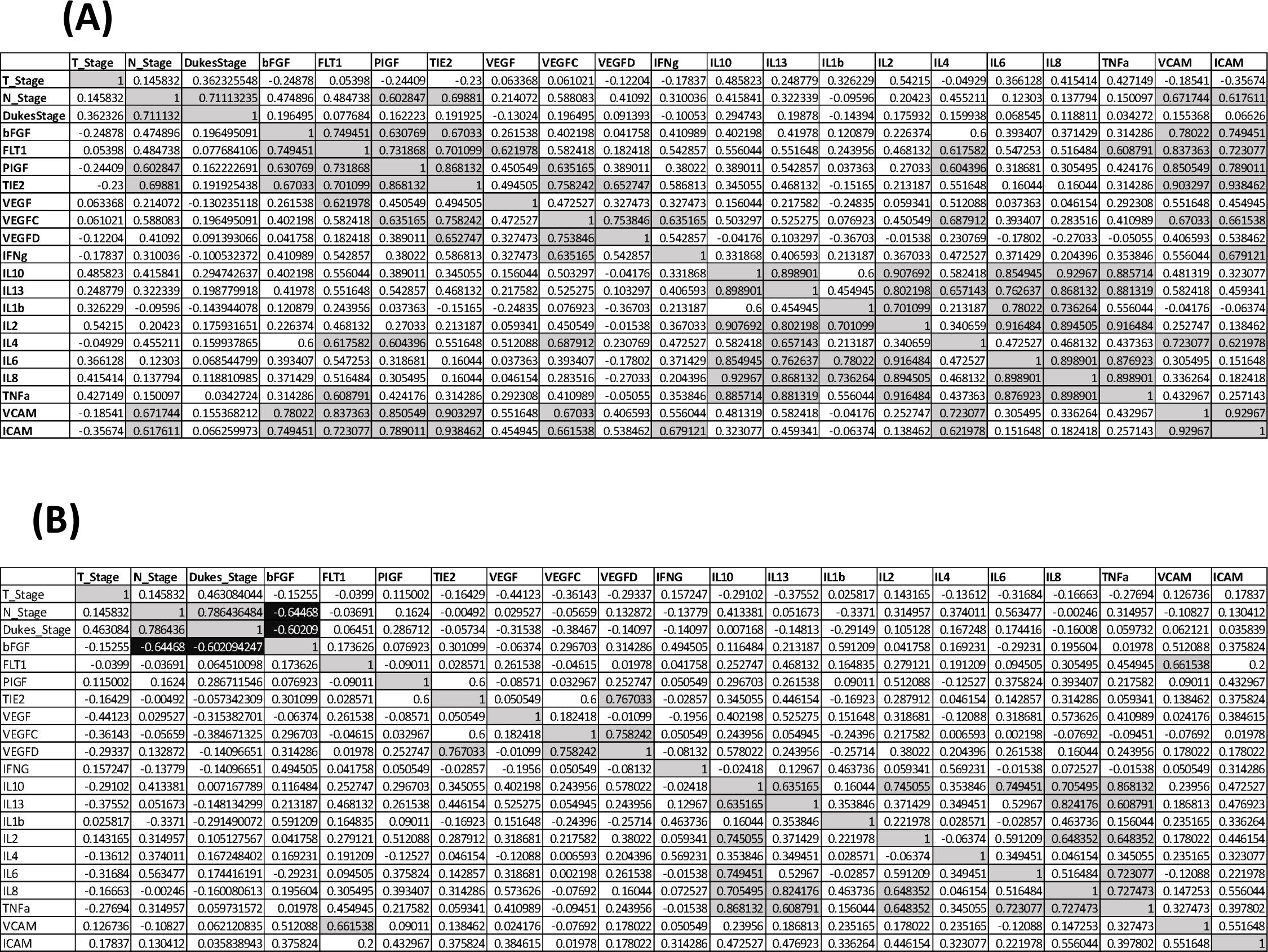
Spearman correlations values clinical characteristics and biological data for n=14 patients. **(A)** Baseline secreted angiogenic and inflammatory analyte levels compared **(B)** Following the addition of Q8 (10uM), the expression of each secreted analyte was compared to its baseline (control) expression level to determine the relative fold change. Quadrants above and below the diagonal are mirror images. Correlation values >0.6 highlighted in grey, correlation values <-0.6 are highlighted in black.

## Abbreviation

PIGF: Placenta Growth Factor
VEGF: Vascular Endothelial Growth Factor
bFGF: Basic Fibroblast Growth Factor
FLT1: Vascular Endothelial Growth Factor Receptor 1
IFNγ: Interferon gamma,
TNFα: Tumour Necrosis Factor alpha

